# Directed differentiation of human pluripotent stem cells into articular cartilage reveals effects caused by absence of *WISP3*, the gene responsible for Progressive Pseudorheumatoid Arthropathy of Childhood

**DOI:** 10.1101/2023.04.01.535214

**Authors:** Chaochang Li, Mireia Alemany Ribes, Rosanne Raftery, Uzochi Nwoko, Matthew L. Warman, April M. Craft

## Abstract

**Objectives:** Progressive Pseudorheumatoid Arthropathy of Childhood (PPAC), caused by deficiency of *WNT1 inducible signaling pathway protein 3* (*WISP3*), has been challenging to study because no animal model of the disease exists and cartilage recovered from affected patients is indistinguishable from common end-stage osteoarthritis. Therefore, to gain insights into why precocious articular cartilage failure occurs in this disease, we made *in vitro* derived articular cartilage using isogenic *WISP3*-deficient and *WISP3*-sufficient human pluripotent stem cells (hPSCs).

**Methods:** We generated articular cartilage-like tissues from induced-(i)PSCs from 2 patients with PPAC and 1 wild-type human embryonic stem cell line in which we knocked out WISP3. We compared these tissues to *in vitro*-derived articular cartilage tissues from 2 isogenic *WISP3*-sufficient control lines using histology, bulk RNA sequencing, single cell RNA sequencing, and *in situ* hybridization.

**Results:** *WISP3*-deficient and *WISP3*-sufficient hPSCs both differentiated into articular cartilage-like tissues that appeared histologically similar. However, the transcriptomes of *WISP3*-deficient tissues differed significantly from *WISP3*-sufficient tissues and pointed to increased TGFβ, TNFα/NFkB, and IL-2/STAT5 signaling and decreased oxidative phosphorylation. Single cell sequencing and *in situ* hybridization revealed that *WISP3*-deficient cartilage contained a significantly higher fraction (∼ 4-fold increase, *p* < 0.001) of superficial zone chondrocytes compared to deeper zone chondrocytes than did *WISP3*-sufficient cartilage. **Conclusions** *WISP3*-deficient and *WISP3*-sufficient hPSCs can be differentiated into articular cartilage-like tissues, but these tissues differ in their transcriptomes and in the relative abundances of chondrocyte sub-types they contain. These findings provide important starting points for *in vivo* studies when an animal model of PPAC or presymptomtic patient-derived articular cartilage becomes available.

**KEY MESSAGES:** *What is already known on this topic:*

- Loss-of-function mutations in *WISP3* cause Progressive Pseudorheumatoid Arthropathy of Childhood (PPAC), yet the precise function of *WISP3* in cartilage is unknown due to the absence of cartilage disease *Wisp3* knockout mice and the lack of available PPAC patient cartilage that is not end-stage. Thus, most functional studies of WISP3 have been performed *in vitro* using WISP3 over-expressing cell lines (i.e., not wild-type) and *WISP3*-deficient chondrocytes.

*What this study adds:*

- We describe 3 new *WISP3*-deficient human pluripotent stem cell (hPSC) lines and show they can be differentiated into articular cartilage-like tissue.
- We compare *in vitro*-derived articular cartilage made from *WISP3*-deficient and isogenic *WISP3*- sufficient hPSCs using bulk RNA sequencing, single cell RNA sequencing, and *in situ* hybridization.
- We observe significant differences in the expression of genes previously associated with cartilage formation and homeostasis in the TGFβ, TNFα/NFkB, and IL-2/STAT5 signaling pathways. We also observe that WISP3-deficient cartilage-like tissues contain significantly higher fractions of chondrocytes that express superficial zone transcripts. These data suggest precocious cartilage failure in PPAC is the result of abnormal articular cartilage formation, dysregulated homeostatic signaling, or both.

*How this study might affect research, practice or policy:*

- This study uses *in vitro*-derived articular cartilage to generate hypotheses for why cartilage fails in children with PPAC. This work prioritizes downstream studies to be performed when pre-symptomatic patient-derived cartilage samples or animal model of PPAC becomes available. It is essential to know how WISP3 functions in cartilage to develop therapies that benefit patients with PPAC and other degenerative joint diseases.

## INTRODUCTION

Progressive Pseudorheumatoid Arthropathy of Childhood (PPAC) is a non-inflammatory, painful, debilitating, autosomal recessive genetic skeletal disease caused by loss-of-function mutations in WNT1 inducible signaling pathway protein 3 (*WISP3*, also known as *CCN6*) ^1^, ^2^. Patients with PPAC appear normal in infancy, but by 4 years of age, they are less active than their peers and begin complaining of aching joints ^2-4^. Visually observed enlargement of the children’s inter-phalangeal joints initially suggests an inflammatory disorder, but the joints are not warm, red, or swollen. Radiologically, the phalangeal bones are widened and there are other signs of spondyloepiphyseal dysplasia ^1-4^. Early onset polyarticular cartilage degeneration occurs in all patients with PPAC. Many teenagers and young adults require total knee and total hip replacement, and many older adults require shoulder, elbow, and ankle replacements. At the time of joint replacement surgery, the cartilage from a PPAC joint is indistinguishable from that of an end-stage osteoarthritic joint.

The mechanism by which *WISP3* deficiency causes precocious cartilage failure is unknown. Two independent *WISP3* knockout (KO) mouse strains showed no evidence of precocious cartilage damage ^5^. Global or chondrocyte-specific overexpression of human *WISP3* in mice also produced no detectable phenotype ^6,7^. Absence of a *WISP3*-deficiency phenotype in mice and lack of pre-end-stage cartilage specimens from patients necessitated studies of *WISP3* function being performed *in vitro*. These studies suggested that *WISP3*, which is a secreted protein and member of the connective tissue growth factor, cysteine-rich protein 61, nephroblastoma overexpressed (CCN) protein family, could regulate multiple signaling pathways including BMP, TGFβ, WNT, p38, integrin, and IGF1 signaling, affect matrix synthesis, and alter mitochondrial energy production ^8-26^. However, results from these studies must be cautiously interpreted since most examined the effects of overexpressing *WISP3* or knocking down *WISP3* in cells other than chondrocytes.

We assessed the effect of *WISP3* deficiency histologically and transcriptionally in cartilage we produced via directed differentiation of human pluripotent stem cells (hPSCs). We previously reported methods to generate chondrocyte lineages that produce articular cartilage-like and growth plate cartilage-like tissues from hPSCs ^27-29^. To determine whether and how *WISP3* deficiency affects this process, we used isogenic hPSCs (induced pluripotent and embryonic stem cells) that have bi-allelic or heterozygous *WISP3* loss-of-function mutations. Bi-allelic loss-of-function mutations cause PPAC, whereas heterozygous loss-of-function mutations are found in asymptomatic carriers. Therefore, we consider hPSCs with bi-allelic mutations to be *WISP3*-deficient and hPSCs with a heterozygous mutation or 2 wild-type (WT) alleles to be *WISP3*-sufficient. Herein we report that *WISP3*-deficient and *WISP3*-sufficient hPSCs differentiate into chondrocytes and produce articular cartilage-like tissue *in vitro*, but differ in their transcriptomes and in the relative abundances of articular chondrocyte subtypes. Transcriptomic differences occur in several pathways that are important during cartilage formation and homeostasis.

## RESULTS

### Generation of hPSCs that are *WISP3*-deficient or *WISP3*-sufficient

We used skin fibroblasts from 2 unrelated patients with PPAC to generate induced pluripotent stem cells (iPSCs) (**table 1; supplemental figure S1A-C**). iPSCs from one patient (PPAC1) are homozygous for the *WISP3* loss-of-function allele, c.156C>A;p.C52X (NM_198239 and NP_937882.2, respectively). iPSCs from the other patient (PPAC4) are compound heterozygous for c.156C>A;p.C52X and an intronic mutation that affects splicing (g.IVS1-763G>T) (**table 1**). Because asymptomatic carriers of this autosomal recessive disease have 1 working *WISP3* allele, we used CRISPR/Cas9 to revert a mutant allele in PPAC1 to WT and refer to this corrected *WISP3*-sufficient clone as PPAC1-C (**supplemental figure S1D**). We created a 3^rd^ *WISP3*-deficient hPSC line by using CRISPR/Cas9 to produce a large bi-allelic deletion from exons 1 to 4 in the commercially available H9 human embryonic stem cell (hESC) line and refer to this *WISP3*-deficient clone as H9-WISP3KO (**table 1; supplemental figure 2**). In total, 3 *WISP3*-deficient hPSC lines (PPAC1, PPAC4, and H9-WISP3KO) and 2 *WISP3*-sufficient hPSC lines (H9 and PPAC1-C) were studied.

**Table 1.** Human pluripotent stem cell lines generated and used in this study

| Cell Line* | Mutation | Location |
| --- | --- | --- |
| PPAC1-iPSCs | Homozygous for c.156C>A | Exon 2 |
| PPAC1-C-iPSCs | Heterozygous for c.156C>A | Exon 2 |
| PPAC4-iPSCs | Compound heterozygous for c.156C>A and g.IVS1-763G>T | Exon 2 and Intron 1 |
| H9 hESCs | Wild-type | - |
| H9-WISP3KO hESCs | Large genomic deletion of 1189bp | Exon 1 to Exon 4 |

\*PPAC1-iPSC line is isogenic with PPAC1-C-iPSC; H9-WISP3KO hESC line is isogenic with H9 hESC

### All WISP3-deficient hPSCs differentiated into chondrocytes and produced articular cartilage-like tissue

Using our established directed differentiation protocol (**figure 1A**) ^27-29^, we observed the 3 *WISP3*- deficient and 2 *WISP3*-sufficient hPSC lines produced articular cartilage-like tissue (**figure 1B-F**). The tissues exhibited metachromatic toluidine blue staining, contained no hypertrophic chondrocytes (**figure 1B-F, upper panels**), and had collagen fibers oriented parallel to the cartilage surface in the superficial zone and perpendicular to the surface in the deeper zone (**figure 1B-F, lower panels**). These data indicate *WISP3* is not required for *in vitro* chondrocyte differentiation and articular cartilage tissue formation, which is consistent with the normal radiographic appearance of joints in young children with PPAC.

**Figure 1.**
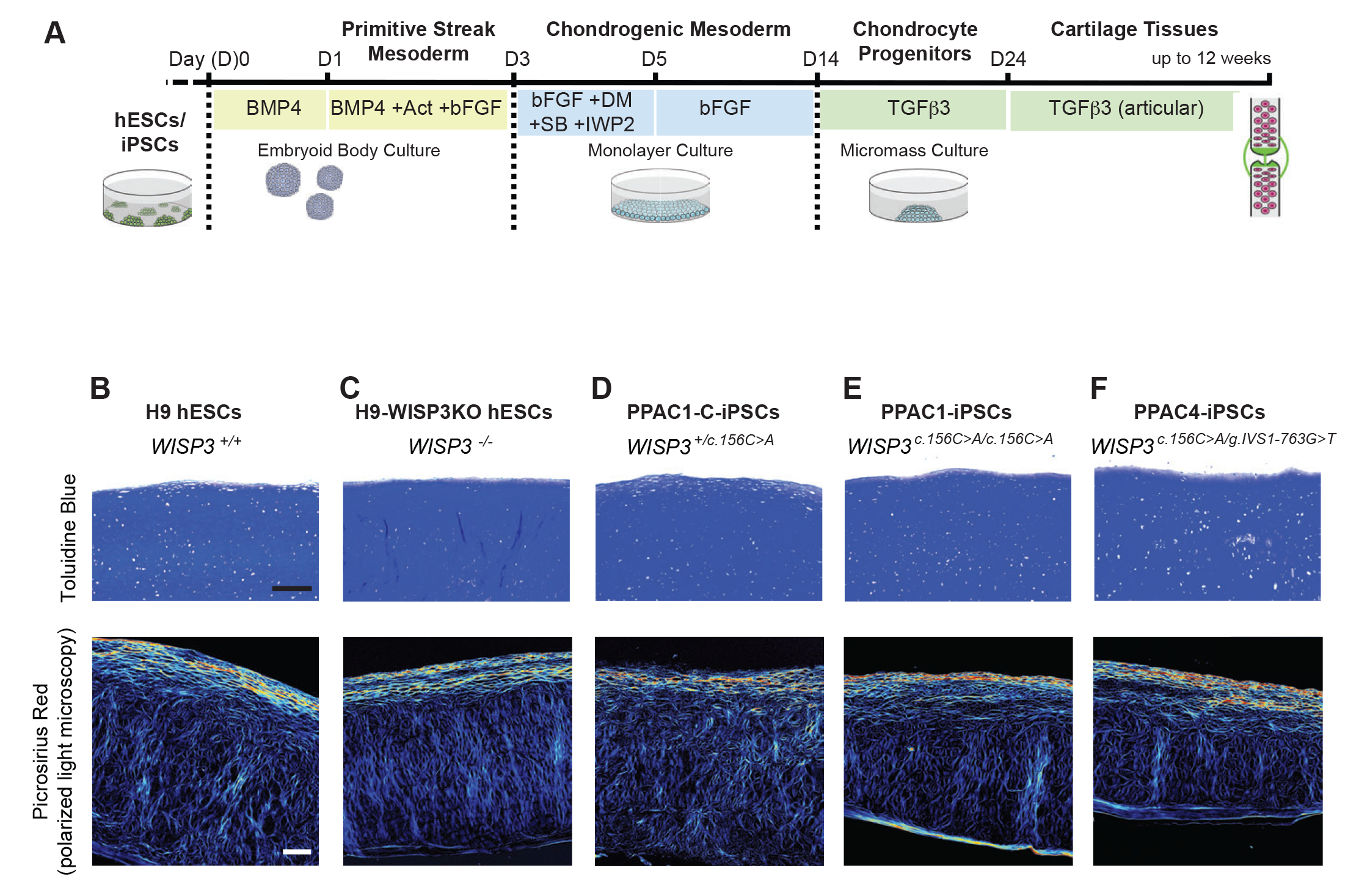
*WISP3*-sufficient and *WISP3*-deficient hPSC lines generate articular cartilage-like tissues *in vitro*. (A) Schematic summary of the directed differentiation method. (B-F) Representative histological images of *in vitro*-derived articular cartilage-like tissues from H9 hESCs (B), H9-WISP3KO hESCs (C), PPAC1-C-iPSCs (D), PPAC1-iPSCs(E) and PPAC4-iPSCs (F). Metachromatic toluidine blue staining (upper panels) indicates sulfated glycosaminoglycans (proteoglycans) are present in the cartilage matrix. Picrosirius red staining (lower panels) coupled with polarized light microscopy depicts the organization of collagen fibers. At the cartilage surface, collagen fibers run parallel with the surface, whereas below the surface the fiber orientation is more perpendicular. Scale bar, 200 µm.

### Bulk RNA sequencing detects transcriptional differences between *in vitro*-derived articular cartilage tissues generated from *WISP3*-deficient and *WISP3*-sufficient hPSCs

To examine the effect of *WISP3*-deficiency on gene expression, we performed bulk RNA sequencing (RNA-seq) on the *in vitro*-derived articular cartilage tissues. Tissues generated from PPAC1-iPSCs and PPAC1-C-iPSCs were tested in triplicate, with RNA sequencing libraries prepared at the same time to minimize batch effects. Principal components analysis (PCA) indicated 58% of the variance in transcript abundance (PC1) was explainable by deficiency of *WISP3*. We observed 2,599 differentially expressed genes (DEGs), with 1,624 and 975 exhibiting increased or decreased mRNA abundance, respectively, in PPAC1- iPSC-derived tissue compared to PPAC1-C-iPSC derived tissue (**figure 2A; supplemental table 4**). Gene Set Enrichment Analysis (GSEA) with these DEGs suggested *WISP3*-defficiency affected several hallmark pathways annotated in the Human Molecular Signatures Database (MSigDB) (**figure 2B**, ranked by p-value).

**Figure 2.**
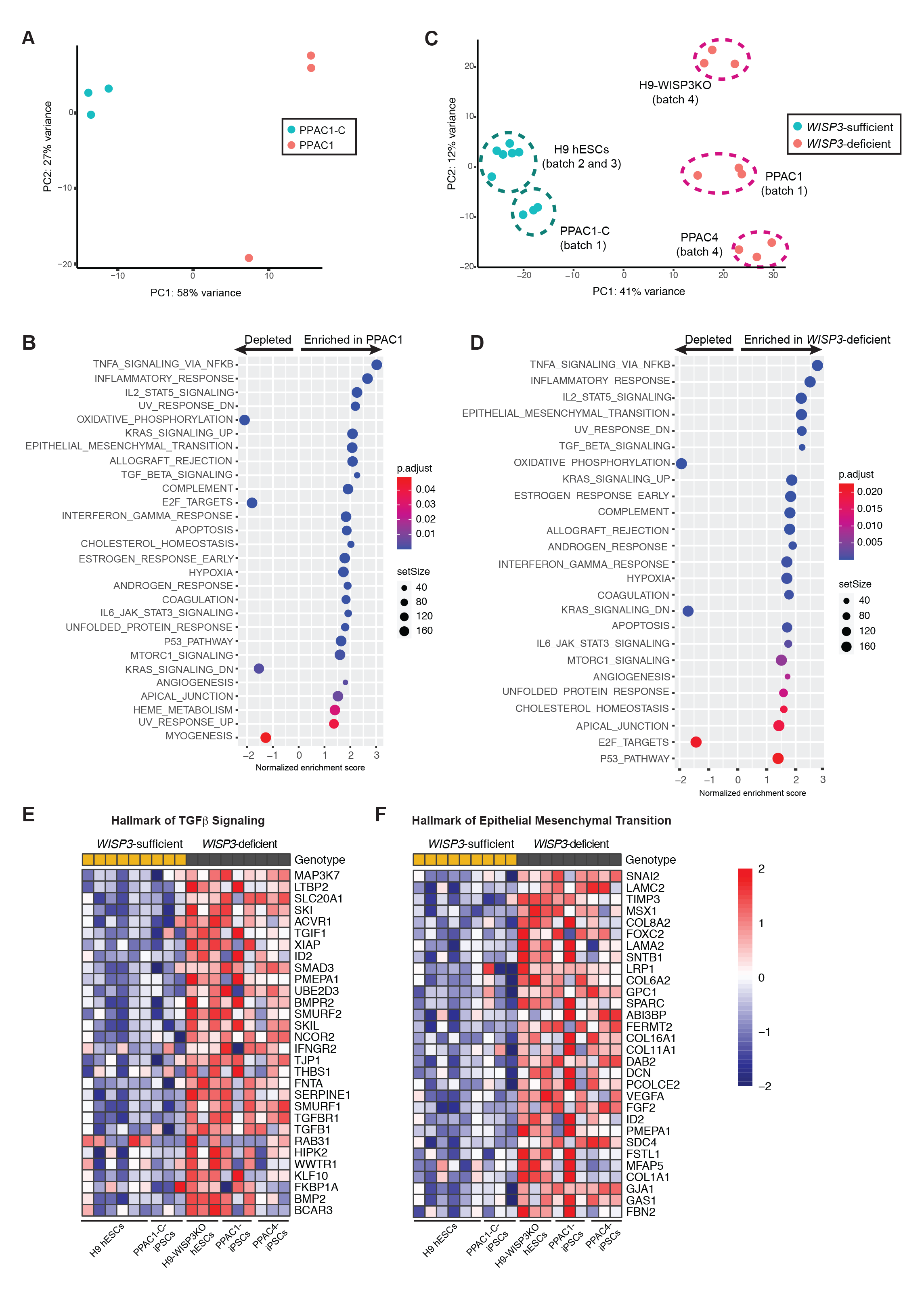
Transcriptomes differ between *WISP3*-sufficient and *WISP3*-deficient hPSC-derived articular cartilage. (A) PCA plot of bulk RNA-seq expression data (batch 1) from PPAC1-C (blue dots) and PPAC1 (red dots) hPSC-derived cartilages. (B) GSEA analysis identifies distinct hallmark pathways significantly enriched or depleted respectively between the *WISP3*-sufficient and *WISP3*-deficient transcriptomes in the batch 1 dataset. (C) PCA plot of all bulk RNA-seq expression data from *WISP3*-sufficient (blue dots) and *WISP3*- deficient (red dots) hPSC-derived cartilages (batches 1 – 4). (D) GSEA analysis using the entire RNA-seq dataset strongly overlaps with that using only batch 1; note that 8 of top 10 GSEA pathways are the same. (E, F) Heatmaps using batch 1 – 4 data, depicting the 30 most significant DEGs (ranked by *p* value) in the TGFβ geneset (E) and in epithelial mesenchymal transition geneset (F). Columns represent biologic triplicates from *WISP3*-sufficient (H9 and PPAC1-C) and *WISP3*-deficient (H9-WISP3KO, PPAC1, and PPAC4) cartilages. Scale is log_2_.

To reduce the number of false positive DEGs and hallmark pathways from the PPAC1-iPSC and PPAC1-C-iPSC comparison, we next included additional bulk RNA-seq data, again performed in triplicate, using *in vitro*-derived articular cartilage tissues differentiated from the H9 hESC, H9-WISP3KO hESC, and PPAC4-iPSC lines. These libraries were created at different times, used different RNA sequencing kits, and were expected to exhibit some batch effects (**supplemental table 1**). When we compared the first dataset (PPAC1-iPSCs vs. PPAC1-C-iPSCs) to this second, mixed genotype dataset, 10 out of the 10 most enriched hallmark pathways remained significant (**supplemental figure S3A**), indicating that similar genes and pathways are affected, regardless of the parental pluripotent stem cell line studied [27 out of 33 total significant pathways remain significant]. Similarly, when we integrated both datasets and compared the transcriptomic changes occurring in all *WISP3*-deficient cartilages, we found that 41% of the variance in transcript abundance (PC1) was accounted by deficiency of *WISP3* (**figure 2C),** and 65% of the initial 2,599 DEGs remaining significant (**supplementary table 5**). Again, 10 out of the 10 most enriched hallmark pathways remained significant (**figure 2B and 2D**) [25 out of 28 total significant pathways remain significant]. Examples of normalized expression changes for genes assigned to two of these hallmark pathways (TGFβ signaling and epithelial to mesenchymal transition) across the panel of *in vitro*-derived articular cartilage tissues are depicted in **figure 2E, F**. Other hallmark pathways genesets are depicted in **supplemental figure S3B-E**.

We orthogonally tested our bulk RNA-seq data by measuring expression levels of a subset of genes in the TGFβ hallmark pathway with qRT-PCR (**figure 3**). We used RNA from the original experiment (**figure 3**, open boxes) and from new experiments (**figure 3**, closed boxes). Cartilage made from cells lacking *WISP3* because of nonsense, frameshift, and splicing mutations lacked measurable *WISP3* mRNA, likely due to nonsense mediated mRNA decay (**figure 3A**). Lubricin (*PRG4*), which is expressed by superficial zone chondrocytes and regulated by TGFβ signaling ^30-33^, was significantly increased in *WISP3*-deficient tissues (**figure 3B**). Other TGFβ responsive genes, connective tissue like growth factor (*CTGF*/*CCN2*) ^34,35^ and the serine protease inhibitor *SERPINE1* ^36^ were also increased in the *WISP3*-deficient tissues (**figure 3C, D**).

**Figure 3.**
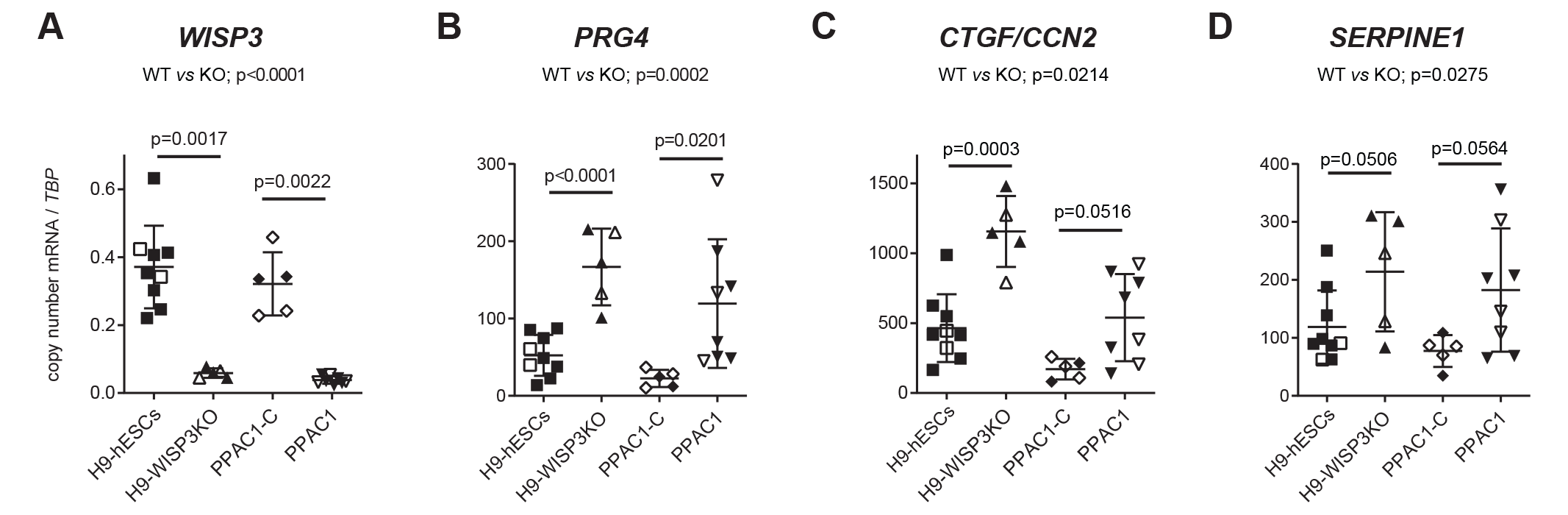
*WISP3* transcripts are absent and TGFβ-responsive transcripts are significantly increased in *WISP3*- deficient *in vitro*-derived articular cartilage tissues. Quantitative RT-PCR values for *WISP3* (A), *PRG4* (B), *CTGF* (C) and *SERPINE1* (D) normalized to *TBP* in H9-hESC-derived (n=9), H9-WISP3KO-hESC-derived (n=5), PPAC1-C-iPSC-derived (n=5) and PPAC1-iPSC-derived (n=8) articular cartilage tissues. Each datapoint represents the mean expression of 2 - 3 tissues from each individual biological replicate experiment/differentiation. Open symbols indicate samples from which bulk RNAseq was also performed; filled symbols indicate new biologic samples. Statistical significance (indicated) calculated by Mann-Whitney U test. Bars show means ± 1 standard deviation.

### Single cell RNA sequencing detects differences in the distribution of chondrocyte subtypes between *WISP3*-deficient and *WISP3*-sufficient *in vitro*-derived articular cartilage tissue

Human articular cartilage is composed of different subtypes of chondrocytes. Superficial zone chondrocytes, which are closest to the cartilage surface, abundantly express *PRG4*. Chondrocytes further from the surface abundantly express chondromodulin (*CNMD*). Increased abundance of a transcript, such as *PRG4*, in bulk RNA-seq data could result from greater expression within each individual superficial cell or a greater fraction of cells within the tissue expressing that transcript. To distinguish between these 2 possibilities, we performed single cell RNA sequencing (scRNA-seq) on the *in vitro*-derived articular cartilage tissues (**figure 4; supplemental figure S4; supplementary tables 2, 7-10**). In the H9 hESC-derived articular cartilage tissues, we identified and named 6 clusters of cells based on their transcriptional profiles and known marker gene expression (**figure 4A; supplemental figure S4A; supplementary table 7**). We detected superficial zone (SZ)-like chondrocytes highly expressing *PRG4* (**figure 4E**) and *elastin* (*ELN*) (**figure 4I**), and intermediate zone (IZ)-like chondrocytes highly expressing *CNMD* (**figure 4M**). We referred to the cluster of cells adjacent to the SZ-like chondrocytes as connective tissue-like, because although they have some uniquely expressed genes (e.g., *tenomodulin*, *TNMD,* **figure 4Q**), they also shared many gene expression patterns with the SZ-like chondrocytes (e.g., *type I collagen* (*COL1A1*) and *PRG4*; **figure 4U; supplementary table 7**). Two additional smaller clusters of cells present in these tissues were named based on their primarily metabolic or proliferation signatures.

**Figure 4.**
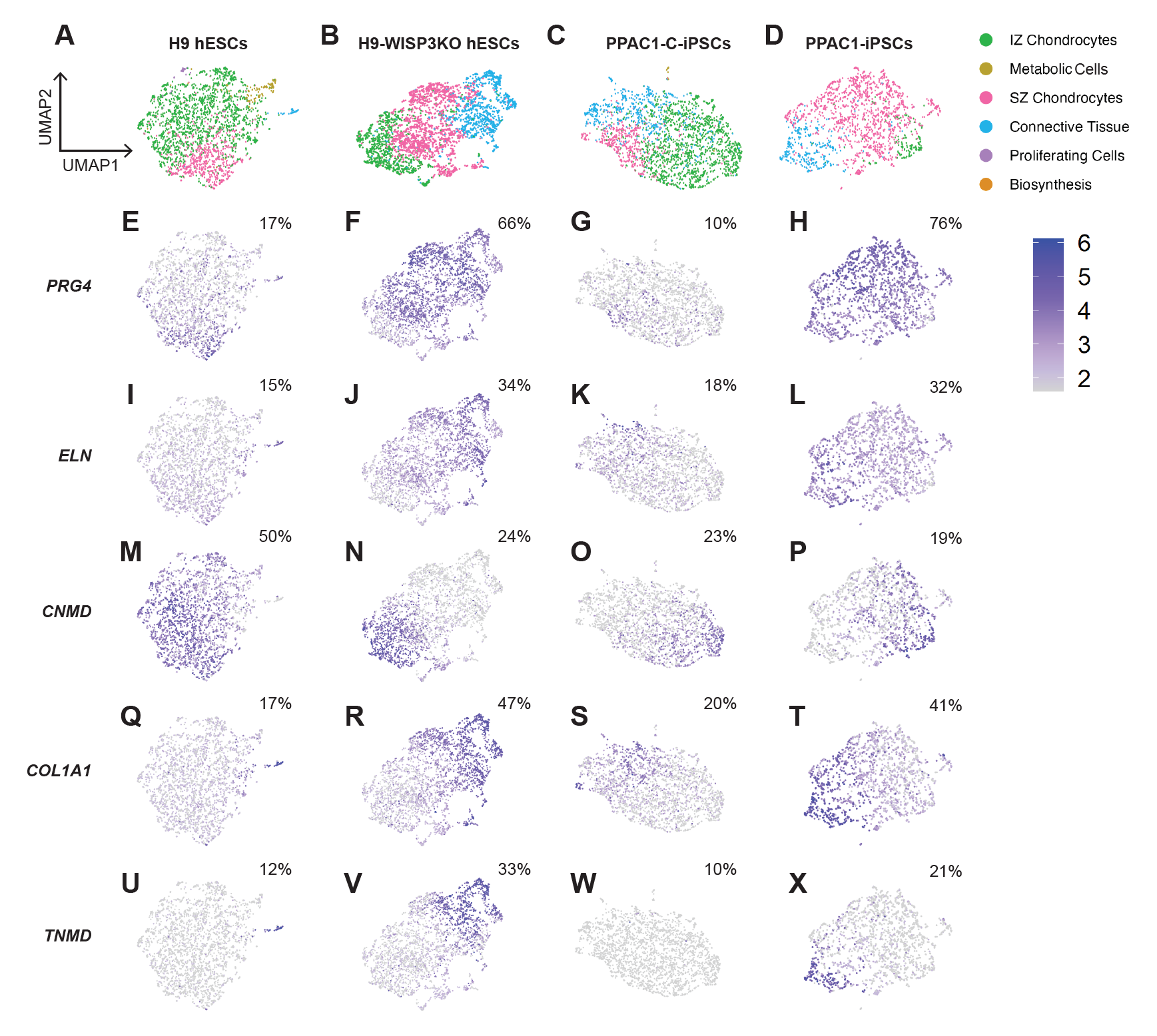
Single cell transcriptomic analyses reveal differences in the cellular composition between isogenic *WISP3*-deficient and *WISP3*-sufficient articular cartilage-like tissues. (A-D) UMAP plots and feature plots (E-P) depicting distinct clusters and gene expression levels within cells isolated from articular cartilage tissues derived from H9 hESCs (A,E,I,M), H9-WISP3KO hESCs (B,F,J,N), PPAC1-C-iPSCs (C,G,K,O) and PPAC1-iPSCs (D,H,L,P) after 12 weeks in culture with TGFβ3. The 6 cell clusters include: SZ chondrocytes, IZ chondrocytes, connective tissue, metabolic cells, proliferating cells and cells with a signature of synthesizing biomolecules. Feature plots illustrate the level of each indicated gene in individual cells (scale is normalized expression, log_2_). The percentage of cells expressing each gene (relative to the total number of cells, using a threshold of normalized counts > 2) within the population is indicated in each panel. SZ chondrocytes highly expressed *PRG4*, and other SZ markers such as *COL1A1*. IZ chondrocytes expressed *CNMD*. Connective tissue/elastic cartilage-like cells expressed *elastin* (*ELN*) and *tenomodulin* (*TNMD*).

When we compared scRNA-seq data between articular cartilage tissues derived from PPAC1-iPSCs and PPAC1-C-iPSCs, and between H9 hESCs and H9-WISP3KO hESCs, transcriptome-based clustering of cell types was similar in *WISP3*-deficient and *WISP3*-sufficient tissues (**figure 4A-D; supplemental figure S4A-D; supplementary table 7-10**). However, *WISP3*-deficient tissues had a larger fraction of cells expressing *PRG4* and other transcripts associated with superficial zone chondrocytes and connective tissue-like cells (**figure 4; supplemental figure S4**), and a smaller fraction of chondrocytes expressing *CNMD*. This was accompanied by increased fractions of cells expressing genes encoding extracellular matrix (ECM) proteins known to sequester growth factors such as TGFβ superfamily ligands in the matrix (e.g., *MFAP5*, *FBN2*; **supplemental figure S4E-L**) and modulate their availability. In addition to observing an increased proportion of *PRG4*-expressing cells after 12 weeks of directed differentiation, we observe a similarly increased fraction after 6 weeks of directed differentiation (**supplemental figure S5; supplementary tables 11, 12**) These data suggest that *WISP3* may determine the relative proportions of chondrocyte subtypes during *in vivo* articular cartilage formation during development.

### *PRG4*-expressing cells are more abundant and extend further beneath the cartilage surface in articular cartilage tissues derived from *WISP3*-deficient hPSCs

We independently confirmed that *in vitro*-derived articular cartilage tissues generated from *WISP3*- deficient hPSC lines contain a larger fraction of *PRG4*-expressing chondrocytes by performing *in situ* hybridization (**figure 5A-B, D-E**). The relative fraction of *PRG4*-expressing cells, and the depth of cartilage below the surface containing them, were higher in *WISP3*-deficient compared to *WISP3*-sufficient tissue (**figure 5C, F**).

**Figure 5.**
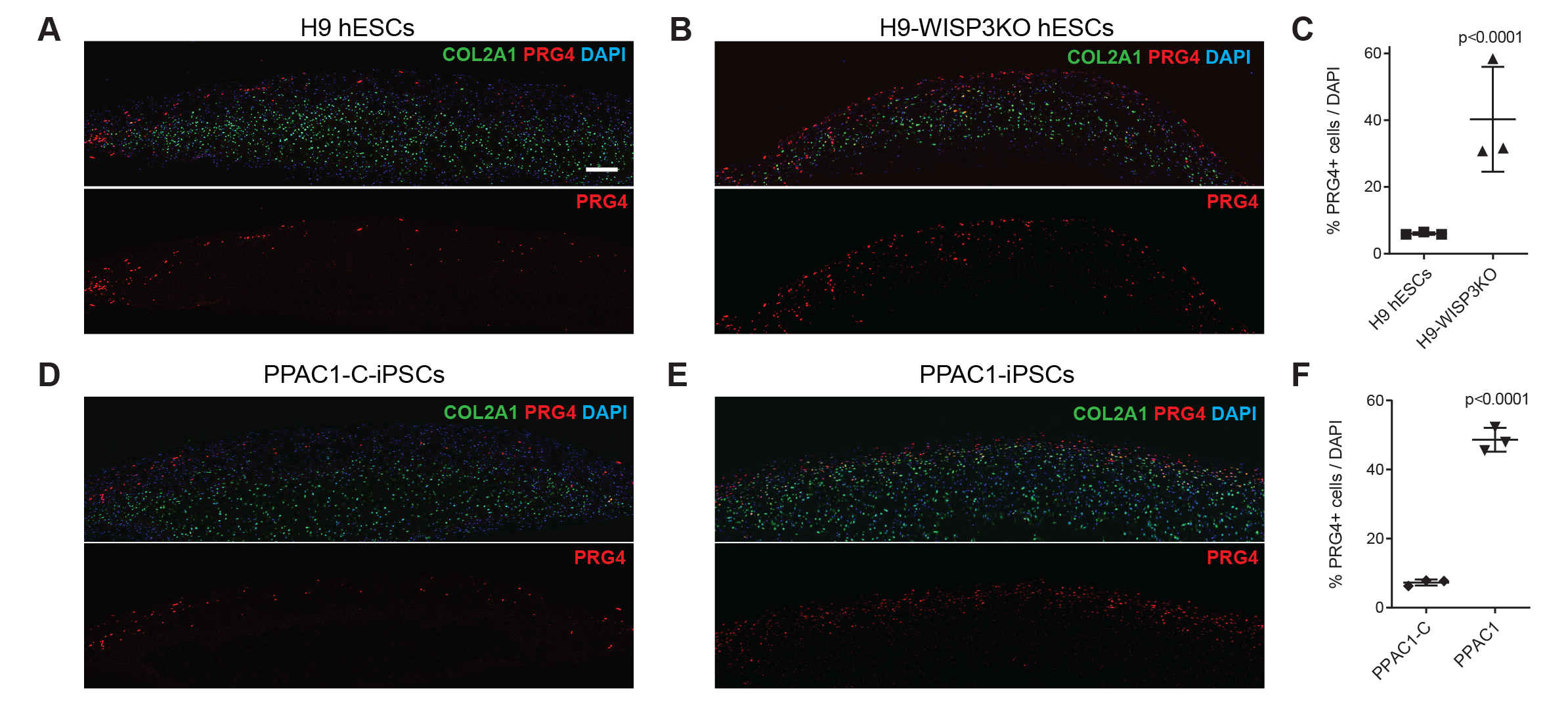
*WISP3*-deficient *in vitro*-derived articular cartilage tissues contain higher percentages of *PRG4*- expressing chondrocytes. (A, B, D, E) *In situ* hybridization for *PRG4* (red) and *COL2A1* (green) within representative sections of *in vitro*-derived articular cartilage tissues derived from H9 hESCs (A), H9-WISP3KO hESCs (B), PPAC1-C-iPSCs (C) and PPAC1-iPSCs (D). Sections were counterstained with DAPI to visualize nuclei. (C, F) The percentage of *PRG4*-expressing cells normalized to the total number of DAPI-positive cells in three representative tissues per cell line was calculated by ImageJ. Scale bar, 200 µm. Data represent three biologically independent tissues. Significance is calculated by student’s t test.

## DISCUSSION

We studied the consequence of *WISP3* deficiency during the directed differentiation of hPSCs into articular cartilage-like tissue. Humans lacking *WISP3* develop precocious cartilage failure, a phenotype that does not occur in *WISP3*-deficient mice ^1,2,5^. Therefore, *in vitro* models that used mouse-derived chondrocytes and other mouse cells may not accurately recapitulate the effect of *WISP3* deficiency in human chondrocytes. Furthermore, there is no study that has examined unaffected or pre-symptomatic cartilage in children with PPAC because of ethical and practical reasons. Therefore, in the absence of a representative animal model and the lack of access to pre-symptomatic patient tissue, we hypothesized that hPSC-derived articular cartilage-like tissues would provide useful insights into the mechanism(s) that contributes to precocious cartilage failure *in vivo*.

We generated 2 PPAC patient-derived iPSC lines and 1 *WISP3* knockout hESC line. We reverted a mutant allele in 1 iPSC line to WT and also used the parental WT hESC line as isogenic controls. Since patients with PPAC appear normal at birth, we anticipated and confirmed that *in vitro*-derived articular cartilage could be generated by *WISP3*-deficient hPSCs (**figure 1**). We then used a combination of bulk RNA-seq, scRNA-seq, and *in situ* hybridization to look for differences between cartilage tissues produced from *WISP3*- deficient versus *WISP3*-sufficient hPSCs. Bulk RNA-seq detected differences in the abundance of ∼ 2000 genes and suggested *WISP3* may modulate several hallmark signaling pathways (**figure 2**).

One such pathway is TGFβ signaling (**figure 2E**). TGFβ signaling is required for cartilage development and homeostasis. In cartilage, *TGFβ1* plays an anabolic role via canonical ALK5/Smad2/3 pathway, and a catabolic role through noncanonical ALK1/Smad1/5/8 pathway ^36,37^. As such, its loss can result in osteoarthritis-like changes in the tissue, and its supplementation can enhance cartilage repair ^38,39^. However, too much TGFβ can damage cartilage; adenoviral-induced over-expression of TGFβ1 in mouse knees caused osteophytes, synovial hyperplasia, and ectopic cartilage-like tissue formation ^40^. Although not performed in chondrocytes, other studies have also implicated *WISP3* in regulating TGFβ signaling *in vitro.* In HME breast cancer cells, shRNA-mediated attenuation of *WISP3* led to up-regulation of *BMP4*, another member of the TGFβ-superfamily ^9^, ^16^. Inhibition of *WISP3* in this same cell line increased mRNA and protein levels of type III TGFβ receptor (*TGFΒR3*), but not *TGFΒR1* or *TGFΒR2* ^17^. Conversely, *WISP3* overexpression in MDA-MB- 231 breast cancer cells reduced *TGFΒR3* protein level *in vitro* and in a xenograft model ^17^. *WISP3* overexpression in zebrafish also appeared to affect BMP signaling, while mutant forms of *WISP3* associated with PPAC did not ^41^. Taken together, these data suggest perturbation of TGFβ signaling contributes to the precocious cartilage failure in patients with PPAC.

It is interesting to speculate how *WISP3* may regulate TGFβ signaling, particularly since we use exogenous TGFβ3 in our directed differentiation protocol to produce articular cartilage-like tissue. WISP3, like other members of the CCN family, contains protein motifs (insulin-like growth factor binding, thrombospondin, von Willebrand factor type C, and cysteine knot domains) that are associated with protein-protein interactions^42^. CCN proteins may therefore affect signaling by sequestering signaling ligands, such as TGFβ from their receptors, or by affecting the efficiency by which a ligand binds to one receptor ^34,42^. There are precedents for each mechanism in other skeletal diseases. For example, failure to adequately sequester BMP when the cystine-knot containing protein *Noggin* is deficient causes multiple synostosis syndrome^43,44^, while mutations that change BMP/BMPR stoichiometry cause defects in chondrogenesis ^45^.

Our *in vitro* data also point to other pathways that *WISP3* deficiency could alter *in vivo*, including the transdifferentiation of cells (e.g., epithelial to mesenchymal transition) and the cellular response to cytokines including tumor necrosis factor, *IL-6* and *IL-2* (**supplemental figure S3**). Hypertrophic chondrocytes and fibrochondrocytes found in osteoarthritic cartilage are postulated to arise from the abnormal differentiation of articular chondrocytes. The cytokine tumor necrosis factor alpha (*TNFα*) is a pro-inflammatory factor and has been implicated in the pathogenesis of osteoarthritis (OA) and rheumatoid arthritis (RA) ^partly^ ^summarized^ ^in^ ^46^. *IL-6*, one of many cytokines present at elevated levels in the synovial fluid of individuals with OA and RA, acts as an activator of the JAK/STAT pathway in both proinflammatory and anti-inflammatory manner ^47^. When synovial fibroblasts were treated with recombinant rat CTGF (rCTGF/CCN2), *p38* MAPK and JNK pathways were increased, and *IL-6* expression was induced due to the increased translocation of *p65*, one of the five components to transport NFkB and c-jun into the nucleus ^48,49^. Interestingly, stimulating human primary chondrocytes or a chondrocyte cell line with TGFβ1 significantly induced *IL-6* mRNA expression ^50^. Finally, GSEA analyses identified a significant depletion of transcripts associated with oxidative phosphorylation in the *in vitro*-derived *WISP3*-deficient tissues (**supplemental figure S3D**) and studies in C-28/I2 and C20A4 chondrocyte cell lines previously suggested *WISP3* also functions intracellularly to stabilize mitochondrial structure and promote oxidative phosphorylation ^20^, ^21^.

We recognize that *in vitro* findings need to be confirmed *in vivo* when cartilage from either a robust animal model of PPAC or a pre-symptomatic patient becomes available. For now, iPSC-based models of articular cartilage may provide the closest *in vitro* platform for identifying pathways that are altered *in vivo*. By starting with iPSCs from affected patients, we are assured that the cells’ genetic background enables the disease-causing *WISP3* mutation to be fully penetrant. Correcting one copy of the mutant allele in these patient-derived cells provides an important isogenic control. Encouraging for studies of *WISP3*, the pathways we identified using *WISP3*-deficient and *WISP3*-sufficient patient-derived iPSCs were also identified using WISP3KO H9 hESCs and their WT counterparts. These data suggest the effect of *WISP3* on chondrocyte differentiation and transcription is large, and not strongly influenced by genetic background or by the type of hPSC used for directed differentiation. The data we obtained using the directed differentiation of hPSCs could be prioritized for *in vivo* study when PPAC tissue becomes available and could be used to test drugs that may ultimately benefit patients.

## ACKNOWLEDGMENTS

The authors would like to acknowledge Dr. Luisa Bonafe and the patients with PPAC and their families who generously donated cells for this study. We thank the Charles H. Hood Foundation (A.M.C.) and the National Institutes of Health and NIAMS (R21-AR076105, A.M.C. and M.L.W.; R01-AR073821, A.M.C) for supporting this work. The authors are grateful to Dr. Laurence Daheron and the Harvard Stem Cell Institute iPSC Core Facility for the generation of iPSC lines and gene-editing services crucial to this work, and to Dr. Luisa Bonafe for critical reading of the manuscript. We also thank the Harvard Medical School Biopolymers Facility, the Bauer Core Facility at Harvard University, and the Center for Skeletal Research Bone RNA-seq and Histology and Histopathology Cores (P30-AR075042) for NGS and histology support.

## DATA AVAILABILITY STATEMENT

Next-generation sequencing data can be accessed in GEO upon publication or is reported previously ^29^.

## AUTHOR CONTRIBUTIONS

All authors were involved in drafting and/or critical review of the manuscript, and approved the final version for submission, taking responsibility for the integrity of the work as a whole. All authors agree to attest to the accuracy and integrity of the work. Conceptualization (A.M.C and M.L.W.), Data Curation (C.L., M.A.R., U.N.), Formal Analysis (C.L. and M.A.R.), Funding Acquisition (A.M.C and M.L.W.), Investigation (C.L., M.A.R., R.R., and U.N.), Methodology (C.L., M.A.R., and R.R.), Resources (M.A.R., R.R., and U.N.), Supervision (A.M.C and M.L.W.), Validation (C.L., M.A.R., and U.N.), Writing – Original Draft Preparation (C.L. and A.M.C.), Writing – Review & Editing (C.L., A.M.C, M.A.R., and M.L.W.).

## METHODS

### Maintenance and differentiation of hPSCs

Maintenance and differentiation protocols have been described in detail previously [27-29]. Briefly, hPSCs (H9- hESCs, H9-WISP3KO hESCs, PPAC1-C iPSCs, PPAC1 iPSCs, and PPAC4 iPSCs) were cultured on irradiated mouse embryonic fibroblasts in DMEM/F12 (Corning) media supplemented with 20% knockout serum replacement (Gibco), nonessential amino acids (Gibco), L-glutamine (Gibco), Pen/Strep (Gibco), β- mercaptoethanol (Gibco), and human bFGF (10 ng/mL) in 6-well tissue culture plates. To differentiate the cells, embryoid bodies (EBs) (containing 3-10 cells) were enzymatically and mechanically generated and cultured in suspension in the presence of BMP4 (1 ng/mL) and ROCK inhibitor (5 µM) for 24 hours in StemPro-34 media (Gibco) supplemented with L-glutamine (Gibco), L-ascorbic acid (Sigma-Aldrich), transferrin (Roche), and a- monothioglycerol (Sigma-Aldrich). On day 1, EBs were harvested and resuspended in StemPro-34 media with bFGF (5 ng/mL), BMP4 (3 ng/mL), Activin A (2 ng/mL), and ROCK inhibitor (5 µM) to induce primitive streak-like mesoderm formation. After 44 hours, on day 3, EBs were harvested and dissociated with TrypLE (Gibco) and cultured as monolayers (100,000 cells per well) in 96-well tissue culture plates (Corning) in StemPro-34 media containing bFGF (20 ng/mL), inhibitor of type I activin receptor-like kinase (ALK) receptors SB431542 (5.4 µM), type I BMPR inhibitor dorsomorphin (4 µM), with or without a Wnt inhibitor IWP2 (2 µM). On day 5, monolayer cultures were then maintained in StemPro-34 media containing bFGF (20 ng/mL) until day 14 to generate chondrogenic mesoderm. From day 0 to day 12, cultures were maintained in 5% O_2_, 5% CO_2_, 90% N_2_ humidified hypoxia environment, and from day 12 forward, cultures were maintained in 5% CO_2_ humidified environment. Articular cartilage-like tissues were generated by plating day 14 chondrogenic mesoderm cells in micromass culture; 250,000 cells were seeded onto 24-well tissue culture plates (Corning) coated with Matrigel (Corning) in base chondrogenic media consisting of high glucose DMEM (Gibco) supplemented with 1% ITS-A, 40 µg/ml L-proline (Sigma-Aldrich), 0.1 µM dexamethasone (Sigma-Aldrich), 100 µg/mL L-ascorbic acid and 10 ng/mL TGFβ3. Micromass tissues were analyzed after 6 and 12 weeks of TGFβ3 treatment.

### Bulk RNA-seq and analysis

Specimen production, RNA isolation and library prep/sequencing were performed in 4 batches based on when articular cartilage-like specimens were differentiated (summarized in supplementary table 1). Cartilage tissues derived from PPAC1-C and PPAC1-iPSCs (Batch 1), H9 hESCs (Batch 3), H9-WISP3KO hESCs and PPAC4- iPSCs (Batch 4), were enzymatically digested with 0.2% type I collagenase (Sigma-Aldrich) for up to 2 hours at 37°C to solubilize the majority of ECM. Liberated cells were pelleted by centrifugation and flash-frozen at - 80°C. RNA was isolated directly from articular cartilage-like tissues derived from H9 hESCs (Batch 2) using trizol.

Total RNA was purified using silica column-based kits (ThermoFisher). RNA quality and quantity were assessed via Bioanalyzer (Agilent), with RNA integrity number (RIN) values > 7. Batch 1 sequencing libraries were prepared by a commercial vendor (Azenta/Genewiz) using the NEBNext Ultra II RNA Library Prep Kit (Illumina) and sequenced using 2 x 150 bp paired end reads. For Batch 2 and 4, libraries were generated by the TruSeq RNA Library Prep Kit v2 (Illumina). For Batch 3, libraries were generated by the KAPA Stranded RNA-Seq Kit (Roche).

Paired-end sequencing reads were mapped to the human reference transcriptome (GRCh37.75) obtained from the ENSEMBL database using the ‘quant’ function of Salmon (version 0.14.0107). Salmon quantification files were imported into R (version 3.6.1) for analysis. For differential expression analysis, a low-count filter was applied to retain the genes that had a quantified transcript count > 5 in at least 3 samples. Statistically significant changes were detected setting the cutoff of *p* value to 0.05 using the ‘DESeq2’ package in R. Multiple functional analyses were performed. The GMT file of gene sets of hallmark pathways was downloaded from the Broad Institute for Molecular Signatures Database (MSigDB). The ‘Combat’ function from the ‘sva’ package was used to minimize effects resulting from batch preparation. Significantly changed hallmark pathways were identified using the ‘GSEA’ function from the ‘fgsea’ package (*p* value < 0.05). Heatmaps of hallmark pathways were generated by the ‘pheatmap’ function from the ‘pheatmap’ package.

### Single-cell RNA-seq

Single cells pooled from 3 articular cartilage-like specimens/cell line, following a 2-hour 0.2% type I collagenase (Sigma-Aldrich) digestion, were individually encapsulated and converted into scRNA-seq libraries according to 10X Genomics Chromium Next GEM Single Cell 3ʹ Reagent Kits v3.1 (10X Genomics). The depth of reads/cell and the number of cells/library for each sample are shown in **supplementary table 2**. We randomly subset 2,716 cells out of 24,792 total cells from PPAC1-iPSCs derived sample for comparisons to PPAC1-C sample. This does not reflect any differences in the ability to extract cells from the tissues using collagenase. The Cell Ranger (10X Genomics) and the ‘Seurat’ package in R (version 3.6.1) were used to assign cells to distinct clusters using principal component analysis which was visualized using UMAP plots. Statistical analysis methods t-Distributed stochastic neighbor embedding (t-SNE) and K-nearest neighbors (Knn) were used to determine the fraction of total cells in each cluster per sample. ‘Seurat’ was used to calculate transcript abundance between cell clusters and between *WISP3*-deficient and *WISP3*-sufficient cells within each cluster.

### Quantitative RT-PCR (qPCR)

Total RNA was extracted from hPSC-derived articular chondrocytes using the MagMAX™-96 Total RNA Isolation Kit (Applied Biosystems). RNA (0.1-1 ug) was reverse-transcribed by Superscript IV VILO reverse transcriptase and treated with ezDNase enzyme (Invitrogen). Quantitative RT-PCR was performed on a ViiA 7 Real-Time PCR System with OptiFlex Optics System (Applied Biosystems) using PowerUp SYBR Green PCR kit (Applied Biosystems). Genomic DNA standards were used to evaluate the efficiency of the PCR and calculate the copy number of each gene relative to the expression of gene encoding TATA-box binding protein (*TBP*). qPCR was performed on cartilage tissues derived from H9 hESCs (n=9), H9-WISP3KO hESCs (n=5), PPAC1-C-iPSCs (n=5) and PPAC1-iPSCs (n=8). Each datapoint represents the averaged expression of 2-3 tissues from each biological replicate (independent experiments/differentiations). Primer sequences are provided in **supplementary table 3**. Mann-Whitney U test was used to detect statistically significant differences between the isogenic cell lines.

### Histology

For toluidine staining, 5 µm sections of hPSC-derived cartilage tissues were deparaffinized by submersion in xylene, rehydrated in a graded series of ethanol solutions, washed in water, stained with 0.1% toluidine blue for 5 minutes, dehydrated and mounted. For picrosirius red staining, 5 µm sections of hPSC-derived cartilage tissues were deparaffinized by submersion in xylene, rehydrated in a graded series of ethanol solutions, washed in water, stained with picrosirius Red Solution for 60 minutes, dehydrated and mounted. 5-8 biological replicates (independent experiments/differentiations) were stained for each cell line. Images acquired on Leica DMi1 inverted microscope. Representative images are shown.

### *In situ* hybridization

The *in situ* hybridization protocol has been previously described [29], and modified for RNAscope as recommended by Advanced Cell Diagnostics (ACD). Briefly, 5 µm sections of formalin-fixed, paraffin-embedded (FFPE) tissues were tested using RNAscope reagents (ACD, Cat. 322381 and Cat. 323110). Target antigen retrieval was performed by incubating sections in TEG buffer (25 mM Tris-HCl pH 8, 10 mM EDTA, and 50 mM Glucose) at 60°C for 4 hours, with multiple buffer changes. Probes utilized were Probe-Hs- PRG4-C3 (ACD, Cat. 427861-C3) and Probe-Hs-COL2A1 (ACD, Cat. 427878). Fluorescence signal was detected using the LSM 800 confocal microscope (Zeiss). The number of cells staining positive for *PRG4* mRNA and the total number of cells staining with DAPI were quantified from 3 biological replicates (independent experiment/differentiation) from each cell line analyzed using ImageJ. Significance is calculated by student’s t test.

### Patient and public involvement

De-identified patient donor cells were obtained for the purpose of this study. Patients and the public were not involved in any steps of the design, conduct, analysis and results dissemination of this study.

---

**Supplemental Figure S1.**
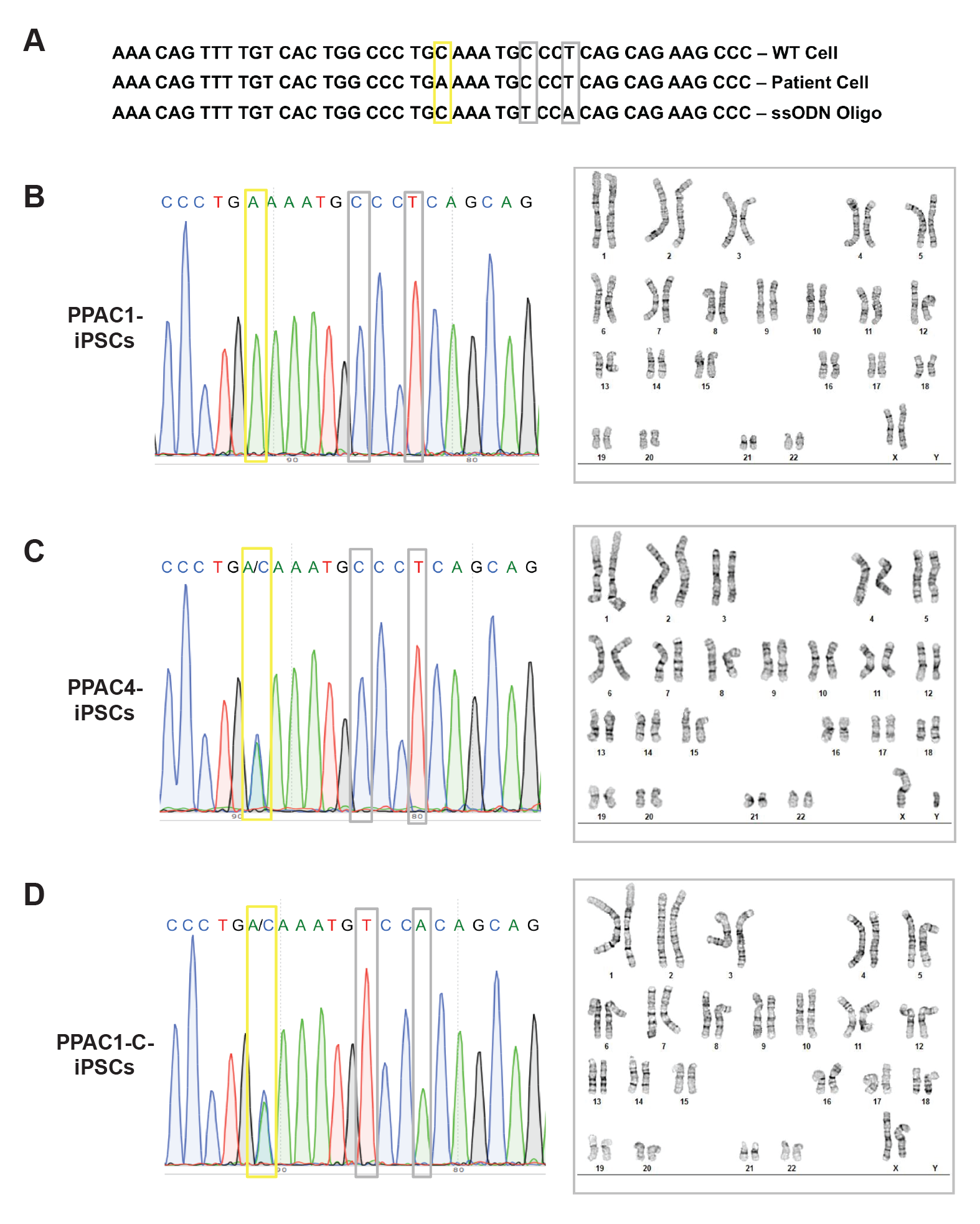
Characterization of PPAC1-, PPAC4- and PPAC1-C-iPSC lines. (A) Sequences of WT and PPAC1 *WISP3* at c.156C>A, along with single-stranded oligo repair template (yellow rectangle); note the repair template also contains 2 synonymous variants (gray rectangles). (B-D) Sanger sequencing results spanning the disease-causing mutation site (left panel) and normal karyotypes of each cell line (right panel). Note the oligo repair template converted the homozygous c.156C>A allele into a heterozygous allele, with both alleles now having the 2 new synonymous variants.

**Supplemental Figure S2.**
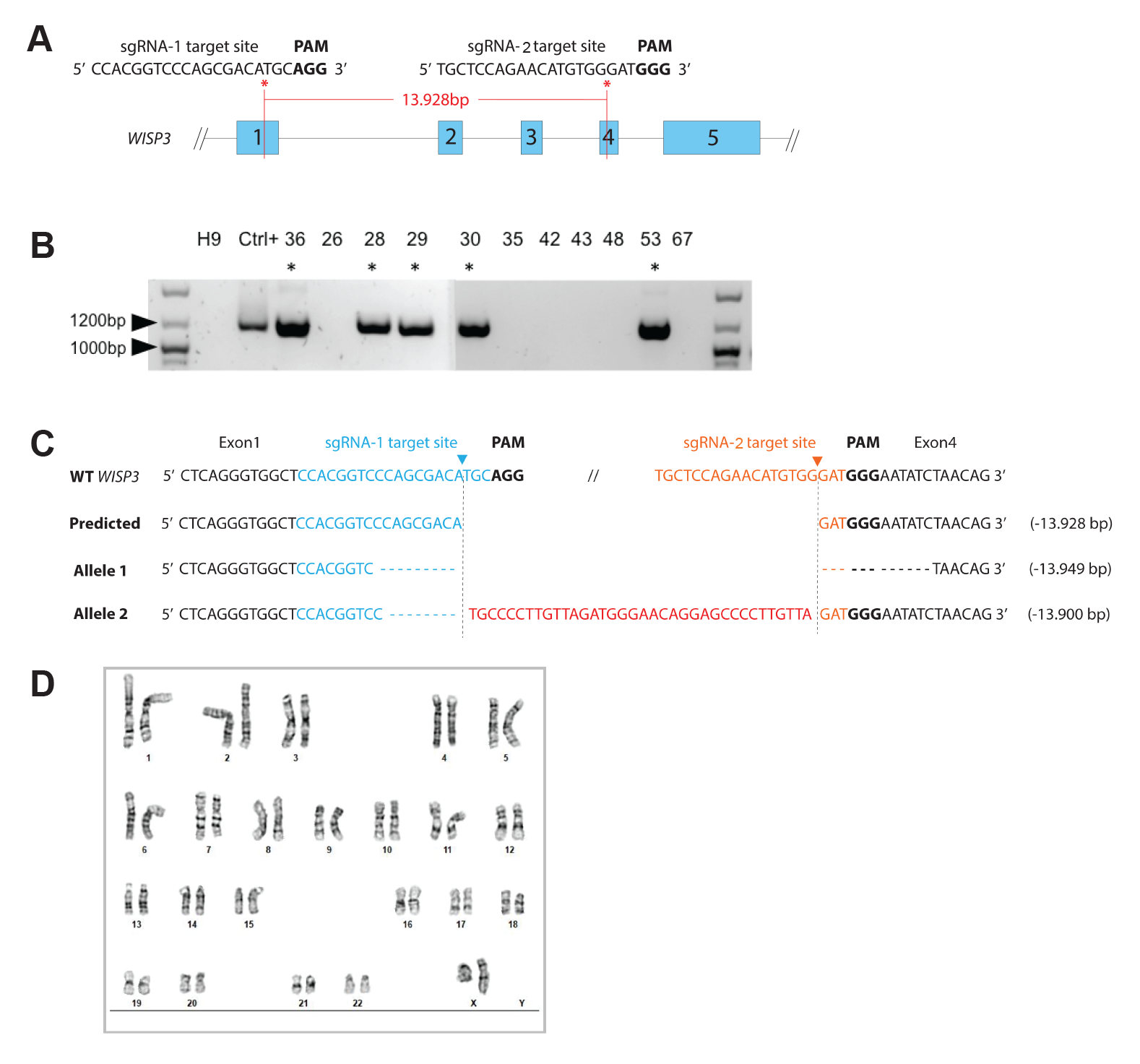
Characterization of the H9-WISP3KO hESC line generated by CRISPR/Cas9 gene-editing methodology. (A) Dual guide RNAs were designed to create a 13,928 bp genomic deletion from exon 1 to exon 4 in the *WISP3* locus. (B) Gel electrophoresis result show PCR products resulting from primers spanning the large genomic deletion in several subclones (sequences in supplementary table 3). The presence of a band indicates at least one allele with a large genomic deletion. (C) Amplicons from clone 36 H9_WISP3KO were subcloned and sequenced, revealing each allele contains a large deletion. Differences between wild-type (WT) allele, a perfect non-homologous end-joining, and the compound heterozygous H9- WISP3KO allele sequences are shown. (D) Normal karyotype of clone 36 H9-WISP3KO hESC line.

**Supplemental Figure S3.**
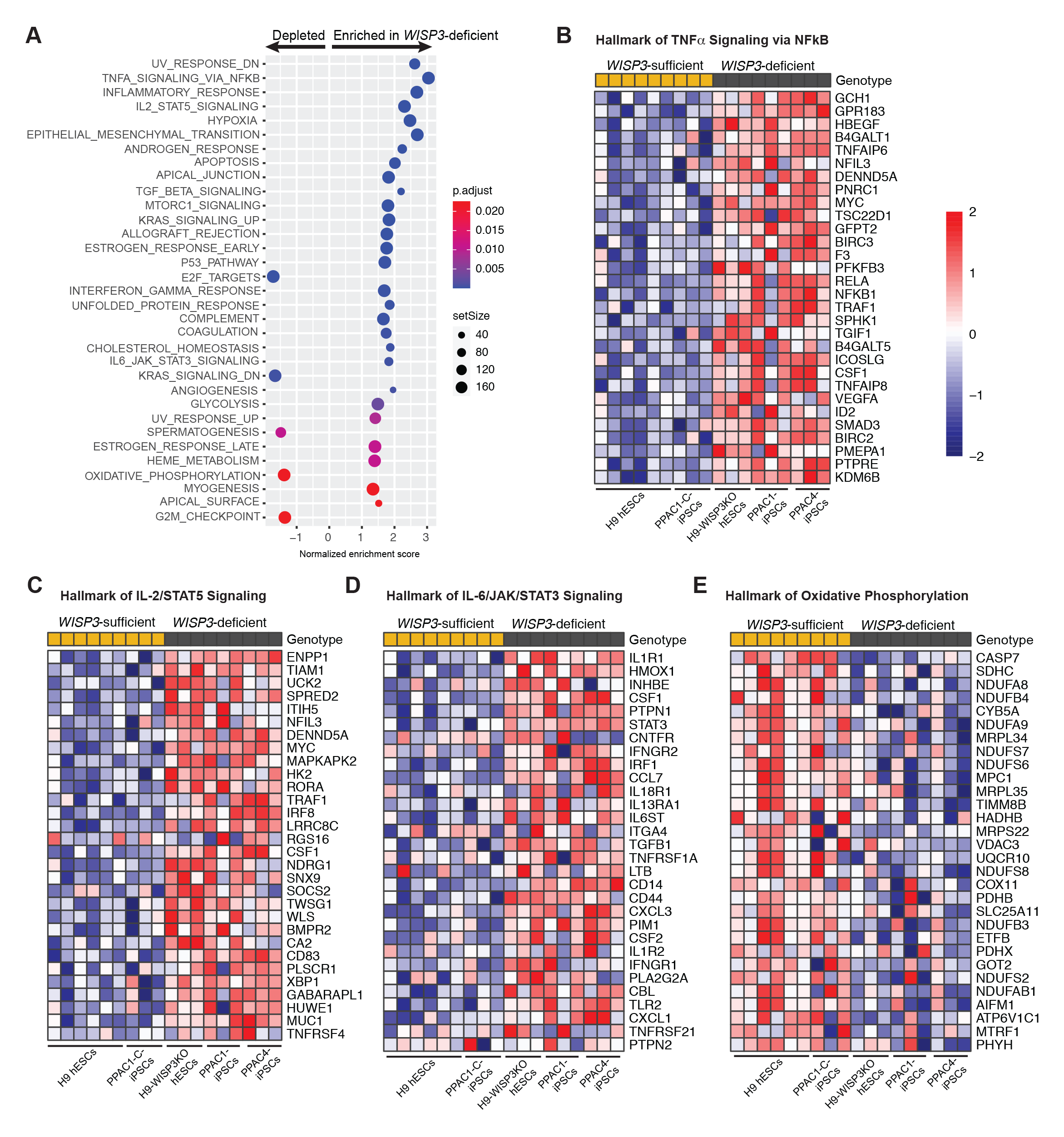
Independent confirmation of hallmark pathway differences between *WISP3*-deficient and *WISP3*-sufficient *in vitro*-derived cartilages. (A) GSEA analysis of bulk RNA sequencing data from H9, H9- WISP3KO and PPAC4 hPSC *in vitro* derived cartilages (batches 2 – 4) strongly overlaps with the batch 1 GSEA analysis; 7 of the top 10 pathways are the same. (B-E) Heatmaps of normalized relative expression levels (log_2_) of the top 30 genes ranked by *p* value in the TNFa/NFkB (B), IL-2/STAT5 signaling (C), IL- 6/JAK/STAT3 (C), and oxidative phosphorylation (D) genesets. Columns represent biologic triplicates with the indicated cell lines.

**Supplemental Figure S4.**
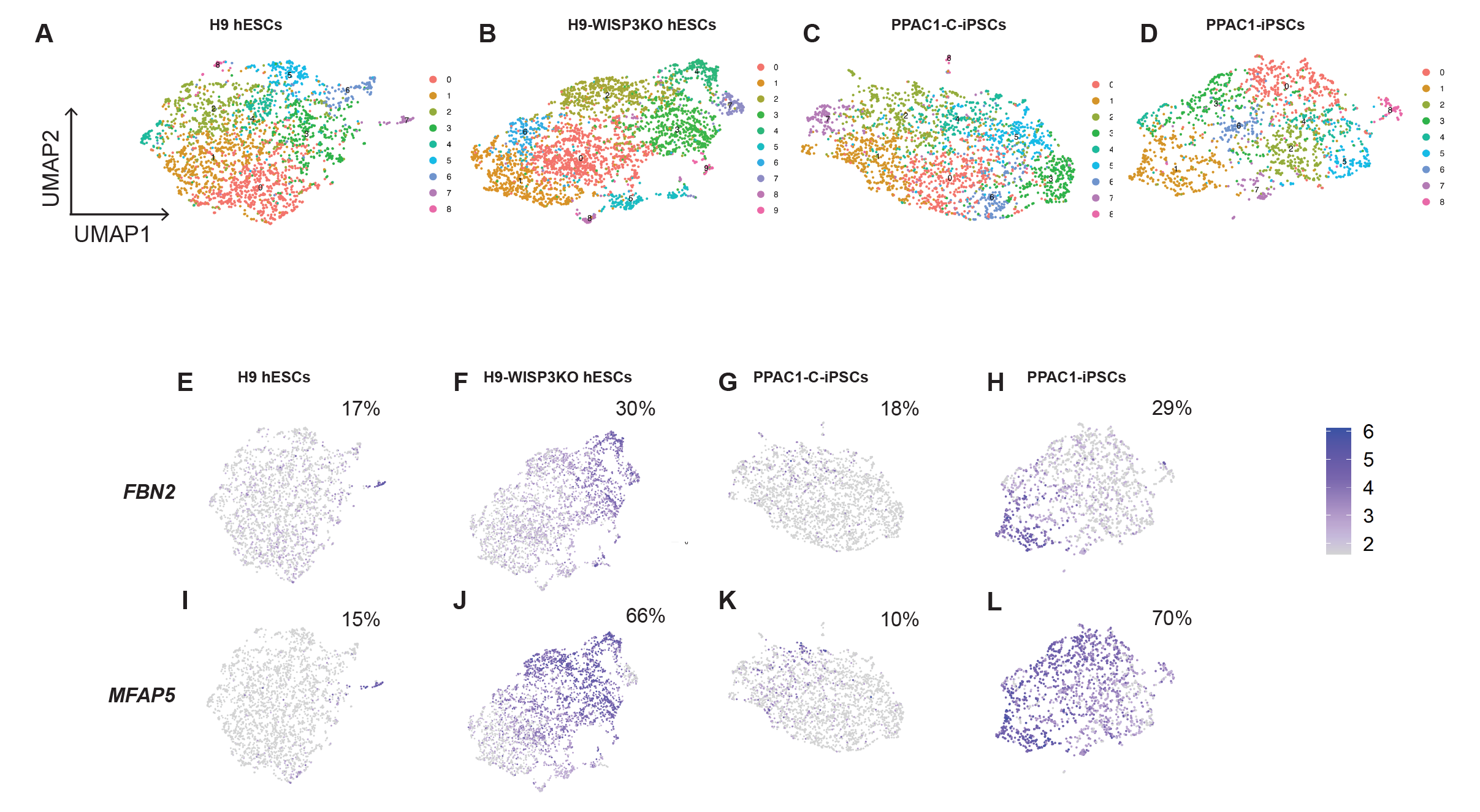
Single cell transcriptomic results for 12-week-old *in vitro*-derived articular cartilage tissues. (A-D) Original UMAP plots depicting all cell clusters before re-naming each cluster based on DEGs and known marker genes of each cell type. Feature plots reveal *FBN2* (E-H) and *MFAP5* (I-L) expression in individual cells. The percentage of cells expressing each gene (using a threshold of normalized counts > 2) within the population is shown on the top right corner in each panel.

**Supplemental Figure S5.**
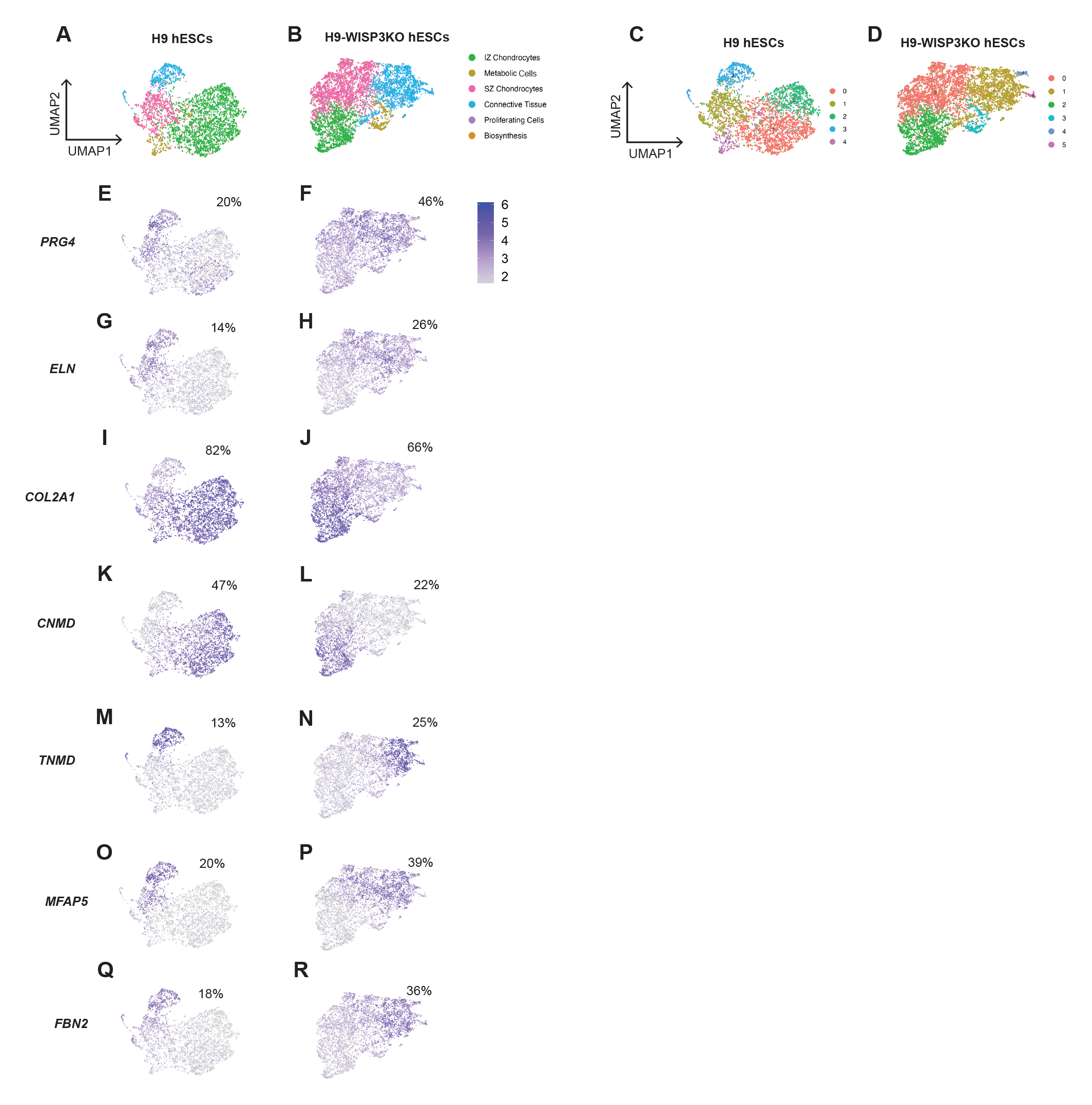
Single cell transcriptomic results for 6-week-old articular cartilage tissues derived from H9 hESCs and H9-WISP3KO hESCs. (A, B) UMAP plots depicting cell cluster identities. (C, D) Original UMAP plot depicting cell clusters determined by Seurat prior to naming. (G-R) Feature plots reveal expression of the indicated genes in individual cells. The percentage of cells expressing each gene (using a threshold of normalized counts > 2) within the population is shown on the top right corner in each panel.

**Supplementary Table 1**

Metadata for bulk RNA-sequencing samples.

**Supplementary Table 2**

Metadata for scRNA-sequencing samples.

**Supplementary Table 3**

Primers used for RT-qPCR.

**Supplementary Table 4**

DEGs between PPAC1-iPSC-derived and PPAC1-C-iPSC-derived articular cartilage tissues.

**Supplementary Table 5**

DEGs between *WISP3*-deficient and *WISP3*-sufficient in vitro-derived articular cartilage tissues.

**Supplementary Table 6**

Normalized counts of genes in *WISP3*-deficient and *WISP3*-sufficient in vitro-derived articular cartilage tissues.

**Supplementary Table 7**

DEGs of each cell cluster identified in H9 hESC-derived articular cartilage tissues.

**Supplementary Table 8**

DEGs of each cell cluster identified in H9-WISP3KO hESC-derived articular cartilage tissues.

**Supplementary Table 9**

DEGs of each cell cluster identified in PPAC1-C-iPSC-derived articular cartilage tissues.

**Supplementary Table 10**

DEGs of each cell cluster identified in PPAC1-iPSC-derived articular cartilage tissues.

